# Upregulation of ATP Citrate Lyase Phosphorylation and Neutral Lipid Synthesis through Viral Growth Factor Signaling during Vaccinia Virus Infection

**DOI:** 10.1101/2023.09.21.558916

**Authors:** Anil Pant, Djamal Brahim Belhaouari, Lara Dsouza, Zhilong Yang

## Abstract

Like all other viruses, poxviruses rely on host cells to provide metabolites and energy. Vaccinia virus (VACV), the prototype poxvirus, induces profound metabolic alterations in host cells. We previously showed that VACV infection increases the tricarboxylic acid (TCA) cycle intermediates, including citrate, that can be transported to the cytosol to be converted to acetyl-CoA for *de novo* fatty acid biosynthesis. ATP citrate lyase (ACLY) is a pivotal enzyme converting citrate to acetyl-CoA. Here, we report that VACV infection stimulates the S455 phosphorylation of ACLY, a post-translational modification that stimulates ACLY activity. We demonstrate that the chemical and genetic inhibition of ACLY severely suppresses VACV replication. Remarkably, we found that virus growth factor (VGF)-induced signaling is essential for the VACV-mediated upregulation of ACLY phosphorylation. Furthermore, the upregulation of ACLY phosphorylation during VACV infection is dependent on the activation of the cellular Akt kinase that phosphorylates ACLY. Finally, we report that VGF-induced ACLY phosphorylation via the EGFR-Akt pathway is important for VACV stimulations of neutral lipid droplet synthesis. These findings identified a previously unknown way of rewiring cell metabolism by a virus and a novel function for VGF in the governance of virus-host interactions through the induction of a key enzyme at the crossroads of the TCA cycle and fatty acid *de novo* biosynthesis. Our study also provides a mechanism for the role played by VGF and its downstream signaling cascades in the modulation of lipid metabolism in VACV-infected cells.

**Importance:** ATP citrate lyase is a key metabolic enzyme that sits at the crossroads of glucose, glutamine, and lipid metabolism. However, how virus infection affects this protein is unclear. Using chemical, genetic, and metabolic approaches we show that VACV, the prototype poxvirus, increases the phosphorylation of ACLY in primary human fibroblasts in a VGF-dependent manner. We further show that the VGF-EGFR-Akt signaling pathway is vital for VACV-induced lipid droplet synthesis. Our findings identified ACLY as a potential target for novel antiviral development against pathogenic poxviruses. Our study also expands the role of growth factor signaling in boosting VACV replication by targeting multiple metabolic pathways.

## Introduction

Poxviruses have significant impacts on public health because of their ability to cause illness and death in both humans and animals. The current global outbreak of mpox, which has been reported in over 110 countries with more than 90,000 cases (including over 30,000 in the USA as of September 1, 2023), highlights the potential for these viruses to cause a pandemic (1). In addition, despite the successful eradication of smallpox, one of the most devastating diseases in human history, there is still a risk of its re-emergence, posing a serious threat to national security (2). On the other hand, many poxviruses are also utilized as vaccine carriers and oncolytic virotherapy agents to combat other diseases (3–5). Vaccinia virus (VACV) serves as the prototype poxvirus and is highly relevant for studying highly pathogenic poxviruses such as mpox and smallpox, given their high genomic similarity with over 95% identical sequences (6).

Metabolism is a battlefront between host cells and viruses during viral infections. The host cell possesses the nutritional resources necessary for viral replication, and viral infections often result in the alteration of the metabolic landscape of the infected cell, with different viruses deploying various strategies to hijack the host cell machinery (7, 8). Although the study of virus-induced metabolic reprogramming has gained considerable interest, the mechanisms underlying the viral repurposing of host cell resources to generate the energy and biomolecules required for viral replication remain largely unexplained. An in-depth understanding of virus-induced metabolic regulation provides ample opportunities for the development of novel antiviral strategies and for uncovering the fundamental mechanisms that regulate cellular metabolism.

VACV induces profound alterations in the host cell metabolism including the TCA cycle. Citrate, which is the first metabolite produced by the TCA cycle, can be shuttled out of the mitochondria to generate acetyl-CoA (9, 10). Acetyl-CoA represents a key precursor of fatty acid biosynthesis and serves as an important source of the acetyl groups for histone acetylation (11). The conversion rate from citrate to acetyl-CoA is governed by the enzyme ATP citrate lyase (ACLY) (10). Therefore, ACLY links carbohydrate metabolism (glycolysis and the TCA cycle), glutamine metabolism (reductive carboxylation), fatty acid synthesis, and histone acetylation making it a pivotal enzyme in cellular metabolism (12, 13). The expression and activity of ACLY are significantly upregulated in several malignancies such as bladder, breast, lung, liver, stomach, prostate, and colon cancers (12, 14–18), and the overexpression of ACLY correlates with poor prognosis in lung adenocarcinoma and blood cancers (15, 19). In addition, the chemical and genetic suppression of ACLY has been shown to inhibit the proliferation and progression of various cancers (20). Because ACLY acts at a critical juncture of host metabolism, ACLY expression levels could be affected by many viruses. However, the mechanisms through which a viral infection may modulate this key host metabolic enzyme and its consequences are lacking. Phosphorylation of ACLY at serine 455 (S455; in humans and mice) also increases enzymatic activity (21). ACLY expression and phosphorylation are regulated by various signals that communicate nutritional status and stimulate growth signaling (20). VACV activates growth factor signaling in infected cells via a VACV-encoded growth factor (VGF), the viral homolog of cellular epidermal growth factor (EGF) and transforming growth factor (22–27). In addition to critical functions in the induction of proliferative effects and viral spread, VGF is important for VACV replication in quiescent cells and mouse (28, 29). That most poxviruses encode their own growth factors despite their host cells expressing growth factors suggests that VGF function is vital and is specifically tailored for poxvirus infection. While it is known that growth factors regulate metabolism, the mechanisms, and biological effects are very diverse. The understanding of context-specific roles of growth factors in metabolism is still at its early stage. We previously demonstrated that VGF is a key viral protein that enhances citrate levels in VACV-infected cells by increasing EGFR and MAPK activation and inducing the non-canonical phosphorylation STAT3 (30). Interestingly, the activation of the PI3K/Akt pathway by EGFR and insulin signaling represents a primary regulatory mechanism for ACLY phosphorylation and activation (20, 21, 31). VACV infection also enhances the phosphorylation of Akt in a VGF-dependent manner (22, 32), prompting us to hypothesize that the VGF-induced PI3K/Akt pathway regulates ACLY phosphorylation during VACV infection.

Our current study reports that VACV infection stimulates the S455 phosphorylation of ACLY, and chemical and genetic inhibition of ACLY severely suppresses VACV replication. VGF-induced growth factor signaling is essential for the VACV-mediated upregulation of ACLY phosphorylation. We further showed that the upregulation of ACLY phosphorylation during VACV infection is dependent on the activation of the cellular Akt pathway. Finally, we report that VGF-induced ACLY phosphorylation via the EGFR-Akt pathway is important for the synthesis of lipid droplets, which are rich in neutral lipids and play an important role in the storage of lipids as the source to fuel the TCA cycle for energy production. These findings identified a novel function for VGF in the governance of virus-host interactions through the induction of a key enzyme associated with host fatty acid metabolism. Our study also provides the mechanism by which VGF and its downstream signaling cascades modulate lipid metabolism during VACV infection. Furthermore, our findings expand the understanding of the role played by growth factors in the regulation of cellular metabolism during a viral infection.

## Materials and methods

### Cells and viruses

Primary HFFs were a kind gift from Dr. Nicholas Wallace at Kansas State University. Primary HFFs, HeLa cells (ATCC CCL-2), and A549 (ATCC CCL-185) were grown in Dulbecco’s modified Eagle medium (DMEM; Fisher Scientific), supplemented with 10% fetal bovine serum (FBS; Peak Serum), 2 mM glutamine (VWR), 100 U/ml of penicillin, and 100 μg/ml streptomycin (VWR) in a humidified incubator at 37 °C with 5% CO_2_. BS-C-1 cells (ATCC CCL-26), and HEPG2 cells (from Dr. Annie Newell-Fugate at Texas A&M University) were cultured in Eagle’s minimal essential medium (EMEM; Fisher Scientific) using the same supplements and environments described for HFF culture. The WR strain of VACV (ATCC VR-1354) was amplified, purified, and quantified using previously described titration methods (33). When the cells reached the desired confluency of approximately 90-95%, they were infected with the indicated MOI of the indicated viruses in DMEM (Fisher Scientific) lacking glucose, L-glutamine, L-asparagine, sodium pyruvate, and phenol red, which was supplemented with 2% dialyzed FBS (Gibco), 100 U/ml of penicillin, and 100 μg/ml streptomycin (VWR). The medium was further supplemented with 1 g/L glucose (Fisher Scientific), glucose plus 2 mM glutamine, or acetate as required. The vΔVGF and vΔVGF_Rev mutant VACVs were generated using a previously described protocol (30).

### Antibodies and chemicals

Antibodies against phospho-ACLY (S455), total ACLY, phospho-Akt (S473), total Akt, perilipin 2, phospho-ACC (S79), total ACC, and horseradish peroxidase-conjugated secondary antibodies were purchased from Cell Signaling Technology. Antibody against the mitochondrial citrate transporter SLC25A1 was purchased from Proteintech. The anti-glyceraldehyde-3-phosphate dehydrogenase (anti-GAPDH) antibody was purchased from Santa Cruz Biotechnology.

The ACLY inhibitor SB 204990, BMS-303141, and NDI-091143 were purchased from Cayman Chemicals. Sodium acetate powder for cell culture was purchased from Sigma-Aldrich. Other chemical inhibitors, including MK-2206 2HCl, H89, stattic, afatinib, and PD0325901 were purchased from Selleck Chemicals and used at the indicated concentrations.

### Cell viability assay

Cell viability assays were performed using a hemocytometer and the trypan blue exclusion assay, as described previously (34). Briefly, after performing each indicated treatment for the indicated time, cells grown in a 12-well plate were harvested with 300 μL trypsin and mixed with 500 μL DMEM using a micropipette. Equal volumes (20 μL) of the cell suspension and 4% trypan blue (VWR) were gently mixed, and the numbers of live and dead cells in each condition were counted using a hemocytometer.

### Western blotting analysis

Western blot was performed as previously described (35). Briefly, after the indicated treatment for the indicated time, the cells were lysed in NP-40 cell lysis buffer and reduced with 100 mM dithiothreitol (DTT), followed by denaturation in sodium dodecyl sulfate-polyacrylamide gel electrophoresis (SDS-PAGE) loading buffer. The samples were boiled at 99 °C for 5 min and separated by SDS-PAGE, followed by transfer to a polyvinylidene difluoride (PVDF) membrane. Membrane blocking was performed for 1 h at room temperature in 5% bovine serum albumin (BSA; VWR) in Tris-buffered saline containing Tween-20 (TBST). The indicated primary antibodies were diluted in the BSA blocking buffer and incubated overnight at 4 °C. After three washes with TBST for 10 min, the membrane was incubated with horseradish peroxidase-conjugated secondary antibody for 1 h at room temperature. Finally, the membranes were developed with Thermo Scientific SuperSignal West Femto Maximum Sensitivity Substrate and imaged using a c300 Chemiluminescent Western Blot Imaging System (Azure Biosystems). If western blotting analysis using another antibody was required, the antibodies were stripped from the membrane by Restore (Thermo Fisher Scientific, Waltham, MA, United States), and the processes of blocking, primary antibody, secondary antibody, and imaging were repeated.

### RNA interference

Specific siRNAs for the indicated target genes and the negative control siRNAs were purchased from Qiagen. The siRNAs were mixed in Lipofectamine RNAiMAX transfection reagent (Fisher Scientific) and transfected to the HFFs in a 6-well plate at a final concentration of 5 nm in OPTIMEM media as per the manufacturer’s instructions. After 48 h, the efficiency of knockdown was confirmed using a western blotting assay.

### Detection of lipid droplets

The cells were seeded on the ibidi 12-well Chamber at a density of 100,000 cells per well. After the cells reached confluency, the cells were washed with PBS and infected with indicated MOI of WT-VACV or vΔVGF in the special DMEM media (Fisher Scientific) with 2% Dialyzed FBS, 2mM glutamine, and 1g/L glucose. After incubation for appropriate time in a humidified incubator at 37°C and 5% CO2 the cells were fixed with 4% paraformaldehyde for 10 minutes at RT. The fixative solution was removed, and the cells were gently washed with 1x PBS buffer 2–3 times to remove residual formaldehyde. The 1000x LipidTOX red neutral lipid stain was diluted 1:1000 in 1X PBS buffer to make a 1x solution and 100uL of the diluted stain was added to each well. The cells were then incubated for 1 h at RT and then washed thrice with 1x PBS before being incubated with 1 μg/mL 2-(4-amidinophenyl)-1H-indole-6-carboxamidine (DAPI) for 20 min at RT in the dark. Finally, the cells were washed three more times with 1x PBS the coverslips, and the silicone gasket were removed, and the culture chamber was mounted on a glass coverslip using ProLong gold mounting medium. The slides were stored in 4°C until confocal microscopy analyses were performed using Zeiss LSM 780 airyscan superresolution confocal microscope. Processing of microscopy images was performed with Zeiss Zen 3.1 (blue edition) and the images were quantified using ImageJ2 (Fiji; version 2.9.0/1.53t).

### Statistical analyses

Unless otherwise stated, the data presented represent the mean of at least three biological replicates. Data analysis was performed in Microsoft Excel (version 16.14) using a two-tailed paired *t-test* to evaluate any significant differences between the two means. The error bars indicate the standard deviation of the experimental replicates. The following convention for symbols was used to indicate statistical significance: ns (not significant), P > 0.05; *, P ≤ 0.05; **, P ≤ 0.01; ***, P ≤ 0.001; ****, P ≤ 0.0001.

## Results

### VACV infection stimulates ACLY phosphorylation

We previously showed that VACV infection increases the levels of citrate (30) and other TCA cycle intermediates in primary human foreskin fibroblasts (HFFs). Citrate can be transported out of the mitochondria into the cytosol, where it is converted to acetyl-CoA and oxaloacetate (OAA), and acetyl-CoA serves as a precursor for fatty acid biosynthesis (13). The conversion of citrate to acetyl-CoA is catalyzed by the enzyme ACLY (**Fig 1A**) (10). The phosphorylation of ACLY at S455 in humans and mice or S454 in rats increases its enzymatic activity (21). Remarkably, we found that VACV infection increased ACLY phosphorylation at S455 in HFFs (**Fig 1B**). To determine the timing of the upregulation of ACLY phosphorylation relative to viral infection, we infected HFFs with VACV and examined ACLY phosphorylation at different times post-infection. We found that VACV infection increased ACLY phosphorylation was observable at 2- and 8-h post-infection (hpi) (**Fig 1C**), indicating that VACV can modulate ACLY activity starting early during the infection.

**Figure 1.**
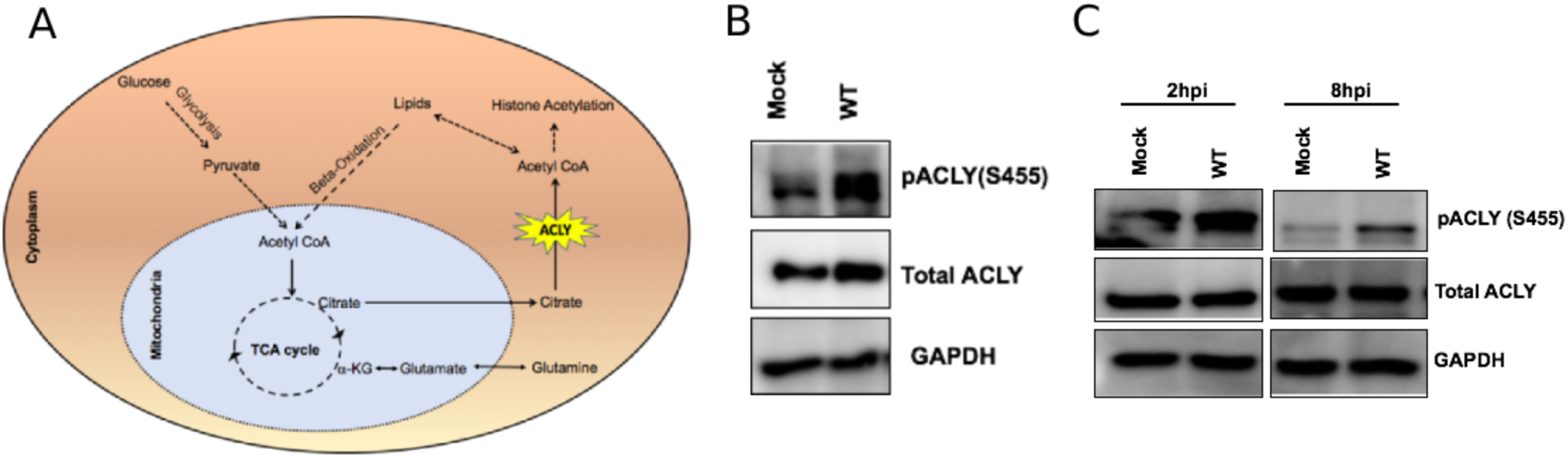
VACV stimulates ACLY phosphorylation. **(A)** ACLY is a key player in cell metabolism. The enzyme ACLY converts citrate, generated by the TCA cycle from glucose or glutamine, into acetyl coenzyme A and oxaloacetate. Acetyl coenzyme A can be further utilized for lipid synthesis, sterol synthesis, and histone acetylation **(B)** VACV infection induces the activation of ACLY phosphorylation at serine 455. HFFs were infected with VACV at a multiplicity of infection (MOI) of 5. Western blotting analysis was performed to measure the levels of ACLY at 4 h post-infection (hpi). **(C)** The upregulation of ACLY S455 phosphorylation can be observed early during VACV infection. HFFs infected with WT VACV at an MOI of 5. The samples were collected at 2 hpi and 8 hpi, followed by Western blotting analysis.

### Inhibition of ACLY suppresses VACV replication

Next, we examined the effects of inhibiting ACLY on VACV replication using SB 204990, a selective and potent inhibitor of ACLY (36). Notably, SB 204990 treatment significantly reduced the VACV titers by 18- and 13-fold in cells infected at a multiplicity of infection (MOI) values of 2 and 0.1, respectively (**Fig 2A**), without affecting the cell viability (**Fig 2B**). To further corroborate these results, we treated the cells with two other sulfonamide-based chemical inhibitors of ACLY, BMS-303141 and NDI-091143 that inhibit ACLY using a different mechanism than that of SB 204990 (37) and assessed the virus replication. BMS-303241 and NDI-091143 reduced the VACV titers by 18 and 16-fold at a high MOI of 2 (**Fig 2C**), and by 3000 and 1428-fold at a low MOI of infection of 0.01 (**Fig 2D).** None of the compounds affected the viability of HFFs at the concentration used in our experiments (**Fig 2E**). Furthermore, we examined the genetic suppression of ACLY levels using small interfering RNAs (siRNAs). ACLY-specific siRNAs effectively reduced the protein expression levels of ACLY (**Fig 2F**) without affecting cell viability (**Fig 2G**). ACLY silencing by two different siRNAs significantly suppressed VACV replication by 10.3 and 8.8-folds (**Fig 2H**). Taken together, these results demonstrated an important role of ACLY activity in VACV replication.

**Figure 2.**
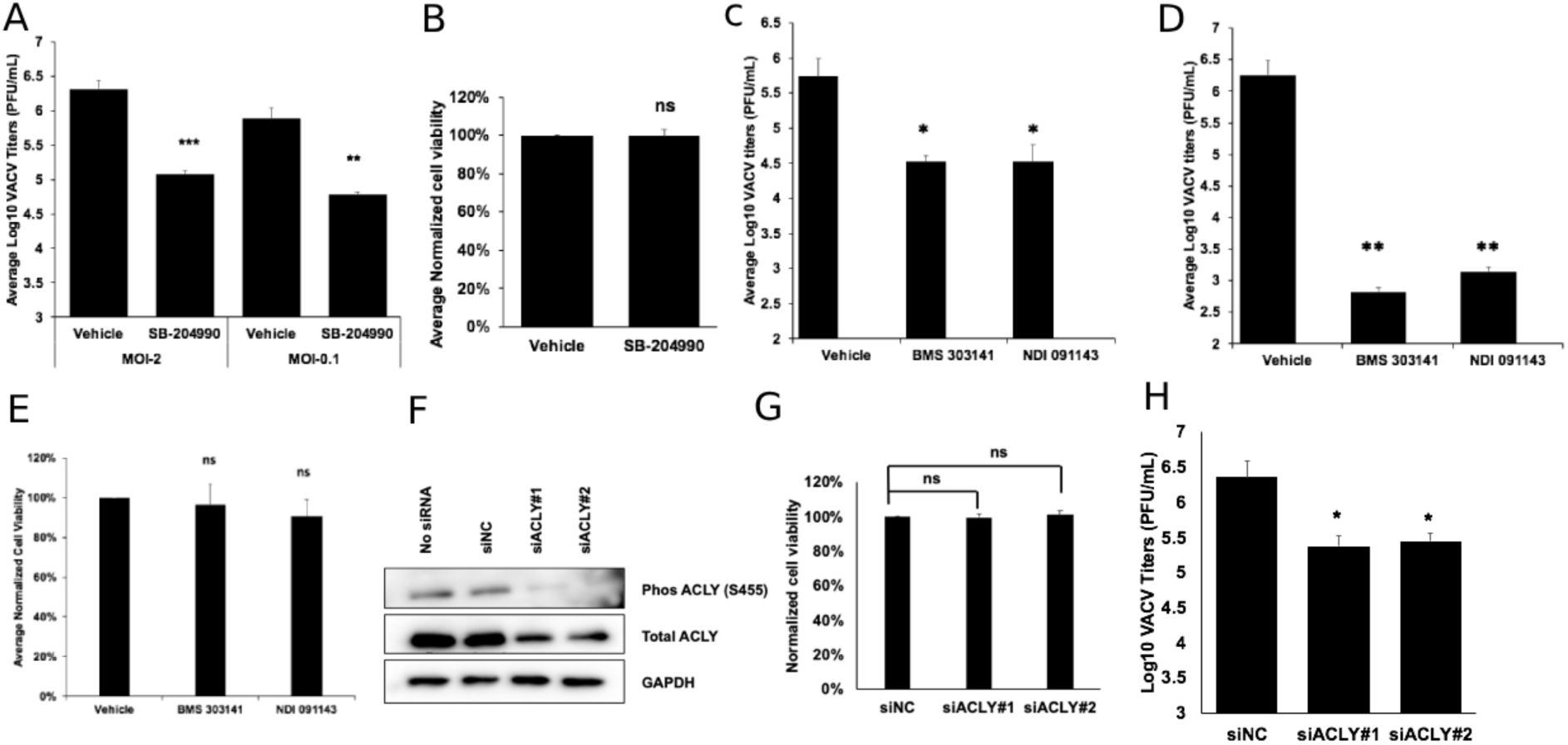
Inhibition of ACLY suppresses VACV replication. **(A)** Chemical inhibition of ACLY suppresses VACV replication. HFFs were infected with VACV in the presence or absence of 100 µM SB-204990. Virus titers were measured by a plaque assay at 24 hpi (MOI = 2) and 48 hpi (MOI = 0.1). **(B)** The inhibition of the ACLY does not alter HFF viability. HFFs were grown in the presence or absence of 100 µM SB-204990 for 48 h. Cell viability was determined by trypan blue exclusion assay using a hemocytometer. **(C, D)** Other chemical inhibitors of ACLY effectively inhibit VACV replication. HFFs were infected with WT VACV in the presence or absence of 10 µM BMS-303141 or 20 μM NDI-091143. Virus titers were measured by a plaque assay at **(C)** 24 hpi (MOI = 2) or **(D)** 48 hpi (MOI = 0.01). **(E)** The ACLY inhibitors do not alter HFF viability. HFFs were grown in the presence or absence of 10 µM BMS-303141 or 20uM NDI-091143 for 48 h. Cell viability was determined by trypan blue exclusion assay using a hemocytometer. **(F)** siRNA-mediated knockdown of ACLY. HFFs were transfected with a negative control siRNA or two specific siRNAs targeting ACLY for 48 h. Western blot was performed to measure the levels of ACLY protein expression. **(G)** ACLY knockdown does not affect HFF viability. HFFs were transfected with the indicated siRNAs for 48 h, and cell viability was determined by trypan blue exclusion assay. **(H)** siRNA-mediated knockdown of ACLY decreases VACV infection. HFFs were transfected with the indicated siRNAs for 48 h and infected with WT VACV at an MOI of 2. Viral titers were measured at 24 hpi. Error bars represent the standard deviation of at least three biological replicates. ns, P > 0.05; *, P ≤ 0.05; **, P ≤ 0.01; ***, P ≤ 0.001.

### Growth factor signaling is essential for the VACV-mediated upregulation of ACLY phosphorylation

Next, we sought to identify the VACV protein responsible for stimulating ACLY phosphorylation. Due to the observed increase in ACLY phosphorylation early during VACV infection, we presumed a viral early protein could be involved. Because VGF is the most highly expressed gene among the 118 VACV early genes (38, 39), and because we previously identified VGF as a key player in the upregulation of citrate levels in VACV-infected cells (30), we tested the role of VGF plays. We used a recombinant VACV from which both copies of the VGF gene were deleted (vΔVGF) from the inverted terminal repeats of the VACV DNA. We also used a VGF revertant VACV (vΔVGF_Rev) by inserting one copy of the VGF gene under its natural promoter but at a different locus in the viral genome (30). We found that infection with vΔVGF abolished this upregulation of ACLY phosphorylation (**Fig 3A**). Notably, phosphorylation could be rescued by infection with vΔVGF_Rev, indicating that VGF is required to induce ACLY phosphorylation (**Fig 3A**).

**Figure 3.**
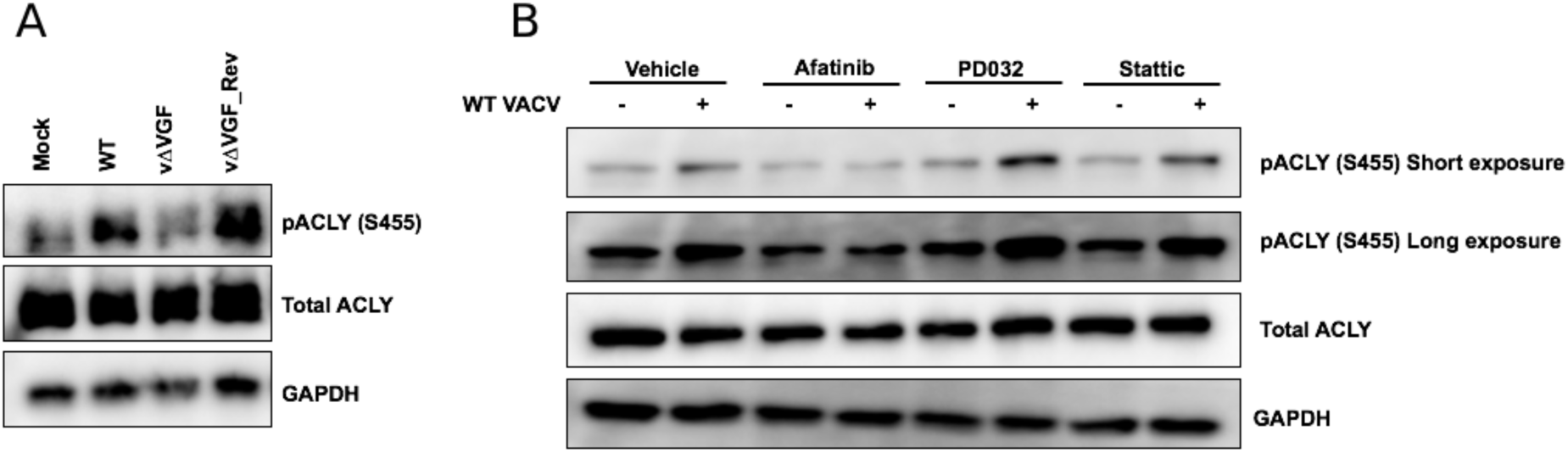
VACV infection induces ACLY S455 phosphorylation in a VGF-dependent manner. **(A)** VGF is crucial for the activation of ACLY phosphorylation (S455). HFFs were infected with the indicated viruses at an MOI of 5. Western blotting analysis was performed to measure the levels of ACLY at 4 h post-infection (hpi). **(B)** VGF-induced epidermal growth factor receptor (EGFR) signaling is required to activate ACLY phosphorylation in VACV-infected cells. Uninfected or HFFs infected with VACV at an MOI of 5 in the presence or absence of 3 μM afatinib, 20 μM PD0325901, or 3 μM stattic were used to detect ACLY levels by Western blotting at 4 hpi.

Because VGF deletion renders VACV unable to increase ACLY phosphorylation, we surmised that VGF-mediated EGFR signaling is involved in the upregulation of ACLY phosphorylation. To explore this possibility, we first tested the effects of an irreversible EGFR inhibitor, afatinib (40), on ACLY levels at a concentration that was previously shown to not affect HFF viability (30). Afatinib treatment reduced the increase in ACLY phosphorylation in VACV-infected cells, with minimal effects observed on uninfected controls (**Fig 3B**). Combined with the previous findings from our lab and others, showing significant reductions in VACV titers following the inhibition of the EGFR pathway (30, 41), our current results indicated that VGF-induced EGFR signaling and ACLY phosphorylation are required for efficient VACV replication.

MAPK is a downstream effector molecule of EGFR (42, 43). We used PD0325901, a selective inhibitor of the MAPK/ERK pathway (43), and found that PD0325901 treatment had minimal effects on ACLY phosphorylation in VACV-infected samples (**Fig 3B**). We have also previously demonstrated that VGF induces the non-canonical phosphorylation of STAT3 at S727 (30). To determine whether STAT3 inhibition affected ACLY phosphorylation, we treated cells with stattic, a well-established STAT3 inhibitor (45). The effects of stattic treatment on ACLY phosphorylation were insignificant (**Fig 3B**). These results suggested that ACLY activation during VACV infection occurs independently of the MAPK and STAT3 signaling pathways.

### VACV infection upregulates ACLY phosphorylation in an Akt signaling-dependent manner

Growth factors activate the PI3K-Akt cascade to elicit a variety of cellular functions (46). Akt is the predominant activator of ACLY phosphorylation (15, 21). Interestingly, VACV infection is known to activate Akt phosphorylation in mouse A31 cells and mouse embryonic fibroblasts, which can be observed at an early post-infection time point (32) and appears to be VGF-dependent in HFFs (22) (**Fig 4A**). We, therefore, examined whether Akt is necessary for the induction of ACLY phosphorylation in VACV-infected HFFs. We measured the levels of ACLY phosphorylation in uninfected and VACV-infected HFFs treated with MK-2206, a highly selective Akt inhibitor (47). MK-2206 treatment reduced ACLY phosphorylation in both uninfected and VACV-infected conditions (**Fig 4B**). The reduction in the uninfected control was less pronounced than that observed in VACV-infected HFFs (**Fig 4B**). In addition to Akt, protein kinase A (PKA) is another well-studied activator of ACLY (48). We used H89, a selective inhibitor of PKA (49), to test the effects of PKA inhibition on ACLY levels following VACV infection. PKA inhibition, however, was not as efficient as Akt inhibition for reducing ACLY phosphorylation in either uninfected or VACV-infected conditions (**Fig 4B**), which suggests that PKA plays a less important role than Akt in ACLY phosphorylation. The chemical inhibition of Akt using MK-2206 also significantly reduced VACV titers by 11- and 21-fold at MOI of 2 and 0.1, respectively (**Fig 4C**), without affecting HFF viability (**Fig 4D**). The findings agree with a previous report showing a reduction of VACV titers upon Akt inhibition in A31 cells and mouse embryonic fibroblasts (32). We further tested the effects of EGFR, MAPK, and STAT3 inhibition on Akt levels during VACV infection. The inhibition of EGFR, but not MAPK or STAT3, suppressed Akt phosphorylation in VACV-infected cells but not in uninfected controls (**Fig 4E**). Taken together, these results indicated that the VGF-induced EGFR pathway serves as an upstream activator of Akt phosphorylation during VACV infection, which results in increased ACLY phosphorylation.

**Figure 4.**
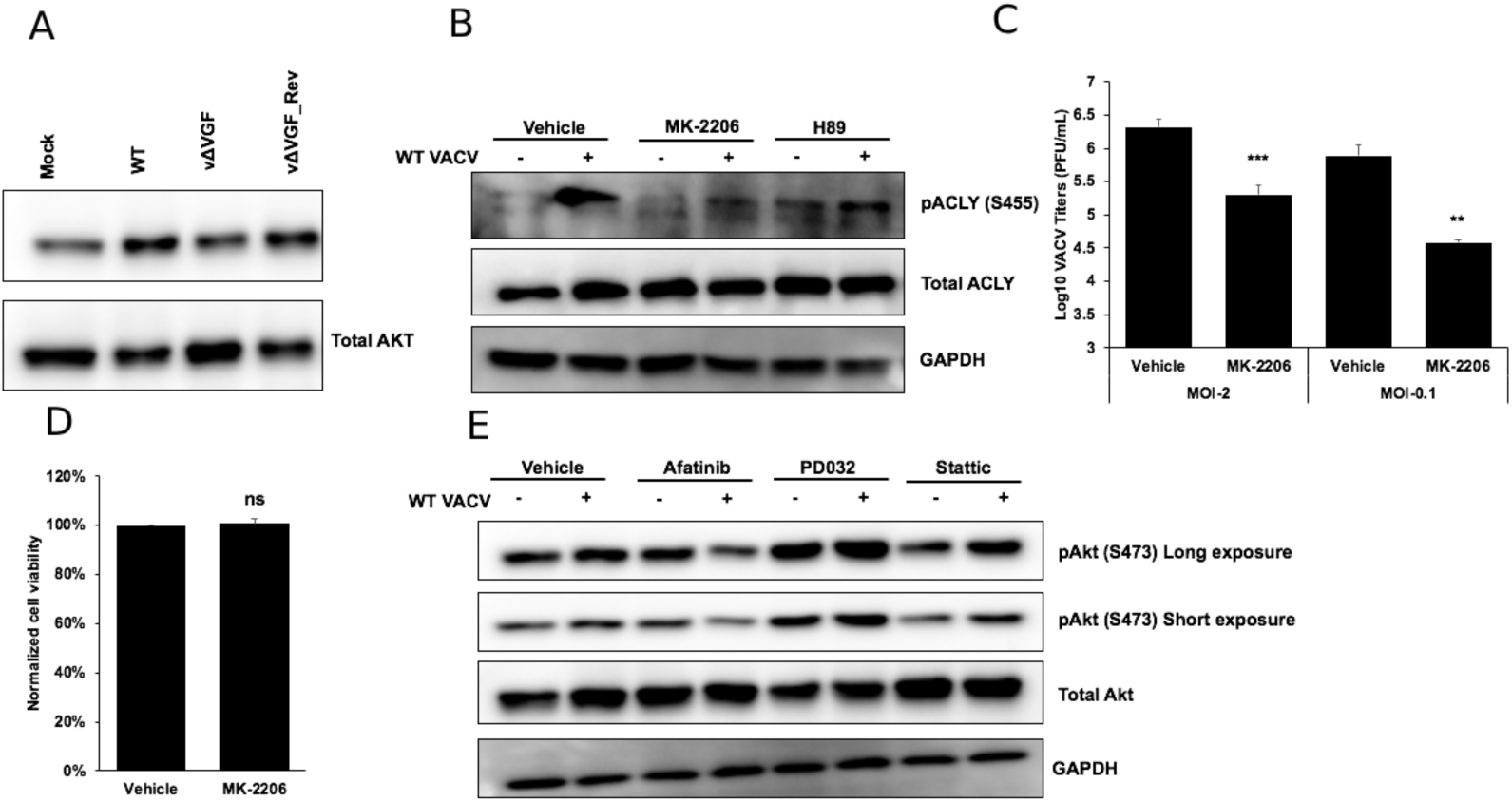
VACV infection upregulates ACLY phosphorylation in an Akt-dependent manner. **(A)** VACV infection induces protein kinase B (Akt) phosphorylation in a VGF-dependent manner. HFFs were infected with the indicated viruses at a multiplicity of infection (MOI) of 5 for 2 h. Western blotting analysis was performed to measure the levels of phosphorylated or total Akt. **(B)** Akt inhibition suppresses ACLY phosphorylation under VACV-infected conditions. HFFs infected with MOI 5 of WT VACV (or uninfected controls) in the presence or absence of 5 µM MK 2206 (Akt inhibitor) or 5 µM H89 (PKA inhibitor) were used to detect ACLY levels by Western blot at 4 hpi. **(C)** The inhibition of the Akt suppresses VACV replication. HFFs were infected with WT VACV in the presence or absence of 5 µM MK 2206. Virus titers were measured by a plaque assay at 24 hpi (MOI = 2) and 48 hpi (MOI = 0.1). **(D)** The inhibition of Akt does not affect HFF viability. HFFs were grown in the presence or absence of 5 µM MK 2206 for 48 h. Cell viability was determined by trypan blue exclusion assay. **(E)** The inhibition of EGFR signaling suppresses Akt phosphorylation upon VACV infection. Uninfected control or HFFs infected with WT VACV at an MOI of 5 in the presence or absence of 3 µM afatinib, 20 µM PD0325901, or 3 µM stattic were used to measure Akt levels by western blotting assay at 4 hpi. ns, P > 0.05; **, P ≤ 0.01; ***, P ≤ 0.001.

### VACV infection induces lipid droplet formation in HFFs through the VGF-EGFR-Akt signaling cascade

A recent study reported that VACV infection increases lipid droplets in mouse bone marrow-derived macrophages (50). Lipid droplets, the cellular organelles that store neutral lipids generated from fatty acids, are a major source of energy production through β-oxidation (51, 52). ACLY predominantly influences the neutral lipid synthesis and mitochondrial β-oxidation in some cancer cells (15, 53). We, therefore, measured the neutral lipids in the VACV-infected HFFs using HCS LipidTOX neutral lipid staining. Interestingly, we found that WT VACV infection substantially increased host cell neutral lipid droplets and vΔVGF infection resulted in a lower level of lipid droplets indicating a vital role of VGF in neutral lipid synthesis (**Fig 5A**).

**Figure 5.**
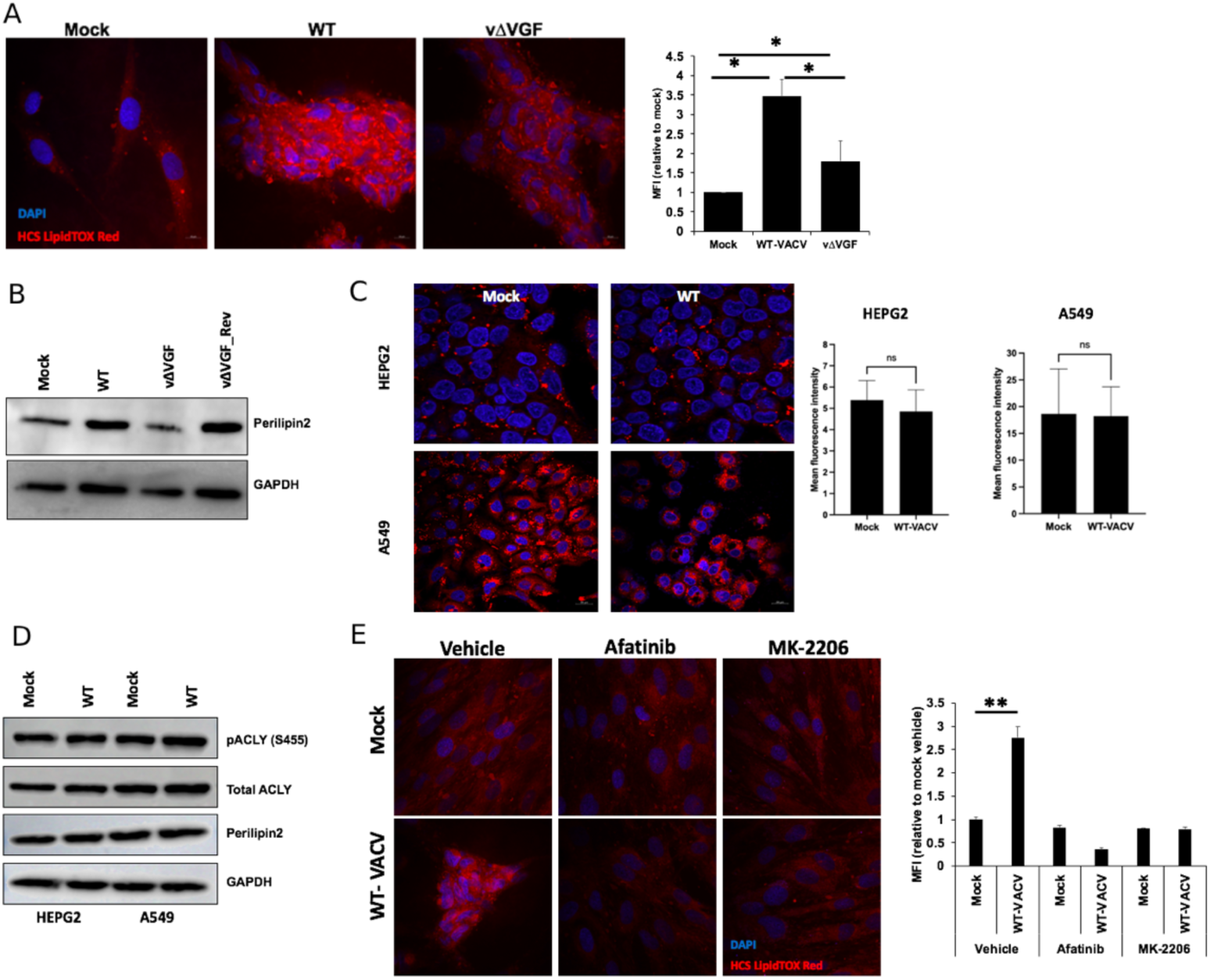
VACV infection stimulates lipid droplet formations in HFFs in a VGF-EGFR-Akt-dependent manner. **(A)** VACV infection induces lipid droplet formation in HFFs in a VGF-dependent manner. HFFs were infected with the indicated viruses at a multiplicity of infection (MOI) of 5 for 8 h. Lipid droplets were stained with HCS Lipidtox Red, and the nuclei were stained with DAPI and imaged under a confocal microscope. The intensity of the red signal that corresponds to the lipid droplets is quantified in the bar graph. **(B)** VACV infection increases the lipid droplet-associated protein levels in HFFs in a VGF-dependent manner. HFFs were infected with the indicated viruses at an MOI of 5 for the indicated 8 h. Western blotting analysis was performed to measure the levels of perilipin 2. **(C)** VACV infection-induced lipid droplet formation is cell-type specific. HEPG2 and A549 cells were infected with VACV at an MOI of 5 for 8 h. Lipid droplets were stained with HCS Lipidtox Red, and the nuclei were stained with DAPI and imaged under a confocal microscope. **(D)** VACV infection does not increase ACLY phosphorylation and lipid droplet-associated protein levels in HEPG2 and A549 cells. HEPG2 and A549 cells were infected with wild-type VACV at a multiplicity of infection (MOI) of 5 for 8 h. Western blotting analysis was performed to measure the levels of ACLY and perilipin 2. **(E)** EGFR and Akt pathways are important for the formation of lipid droplets during VACV infection. HFFs were infected with wildtype VACV at a MOI of 5 for 8 h in the presence or the absence of 3 µM afatinib (EGFR inhibitor) or 5µM MK-2206 (Akt inhibitor). Lipid droplets were stained with HCS Lipidtox Red imaged under a confocal microscope. The intensity of the red signal that corresponds to the lipid droplets is quantified in the bar graph.

Perilipin 2 (PLIN2) is a protein that coats intracellular lipid storage droplets (54, 55). Although traditionally thought to be expressed only in adipocytes, emerging studies have identified an important role of this protein in non-adipocytic cells such as fibroblasts (56). In agreement with the lipid droplet staining, we found that VACV infection increased the levels of PLIN2 in HFFs and this increase was not observed in cells infected with vΔVGF (**Fig 5B)**. Remarkably, PLIN2 levels were rescued by infection with vΔVGF_Rev (**Fig 5B),** indicating that VGF is a crucial viral protein required to form lipid droplets.

We further examined the effect of VACV infection on lipid droplet formation in human hepatoma cell line (HepG2) and human lung cancer cell lines (A549). Unlike in the primary HFFs, we found that VACV infection of HepG2 cells or A549 cells did not increase the HCS LIPIDTOX Red staining (**Fig 5C**). Furthermore, the phosphorylation of ACLY and the levels of PLIN2 also did not increase in either of the two cells following VACV infection (**Fig 5D**) indicating that the neutral droplet formation upon VACV infection could be cell-type and ACLY-dependent. The transformed cells may already have a hyperactive lipid metabolism and may not respond to VACV infection.

Finally, we examined the role of EGFR and PI3K-Akt pathway in neutral lipid droplet formation in VACV-infected HFFs. We found that inhibition of EGFR and PI3K-Akt pathway using specific inhibitors reduced the neutral droplet levels upon VACV infection (**Fig 5E**). Taken together, our results show that VGF-induced ACLY phosphorylation via the EGFR-Akt pathway is important for neutral lipid synthesis and vital for VACV replication.

## Discussion

In the present study, we revealed ACLY that was previously unknown to be targeted by a virus for modulation, is stimulated by VACV infection. We report that VACV infection increases the phosphorylation of ACLY and lipid droplet formation and provide evidence to support the dependence of this activation on VGF, the VACV homolog of cellular EGF. EGFR-induced Akt phosphorylation is critical to the enhancement of ACLY phosphorylation in VACV-infected cells. We further show that VGF and EGFR pathways are important for the formation of neutral lipid droplets that could be converted to fatty acylcarnitines to feed the TCA cycle for bioenergetic requirements (**Fig 6**). Taken together with our previous finding that VGF-induced EGFR activates non-canonical STAT3 phosphorylation to increase TCA cycle intermediate levels (30) these findings present VGF as a versatile protein involved in hijacking host metabolism during VACV infection by modulating different key cellular signaling pathways and different aspects of the host cell metabolism.

**Figure 6.**
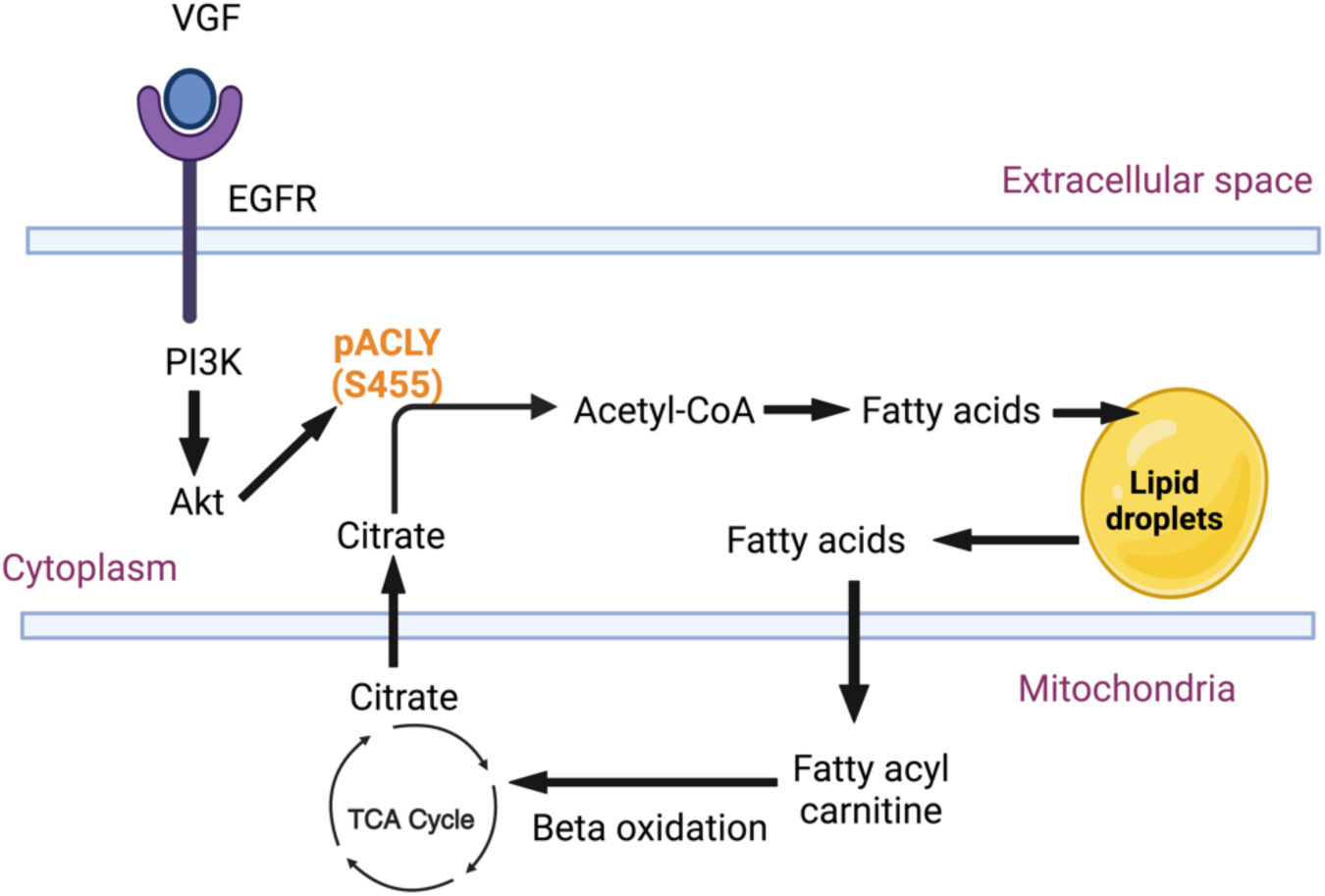
Proposed mechanism and biological impact of induction of lipid droplets during the infection of HFFs with VACV. VACV infection increases the ACLY phosphorylation in a VGF-EGFR-PI3K-Akt dependent fashion that leads to increased neutral lipid droplet formation, geared toward generating β-oxidation intermediates that are eventually recycled to the TCA cycle to generate energy. Created with BioRender.com.

VACV induces profound alterations of the metabolism in its host cells (30, 57–59). VACV induces oxidative phosphorylation (OXPHOS) and the oxygen consumption rate (OCR) indicating the increased energy metabolism during VACV infection (58). Even when most host cell translation is suppressed, VACV induces the selective upregulation of OXPHOS-associated mRNA translation (60). VACV infection enhances the levels of several intermediates of the tricarboxylic acid (TCA) cycle, including citrate (30). Interestingly, VACV also depends on *de novo* fatty acid synthesis to generate an energy-favorable environment (58), indicating the need for VACV to modulate fatty acid metabolism. We previously reported that VACV infection is associated with an increase in the levels of carnitine-conjugated fatty acids, which are essential for β-oxidation, whereas the steady-state levels of long-chain fatty acids were reduced (30), suggesting that the VACV infection-induced changes in fatty acid metabolism. Here, we unveiled the molecular mechanisms underlying the VACV-mediated modulation of a key step linking the TCA cycle and fatty acid metabolism.

VACV infection increases ACLY phosphorylation in a VGF-dependent manner. ACLY sits at the crossroads of the TCA cycle, fatty acid metabolism, and glutamine metabolism (**Fig 1A)**. Interestingly, VACV induces changes in all three aforementioned aspects of cell metabolism, suggesting that ACLY serves as a key regulator in the mediation of VACV-host interactions at the metabolism interface. We have previously shown that VACV infection increases the levels of TCA cycle intermediates (30). Paradoxically, despite the increased phosphorylation of the catalyst (ACLY), VACV infection appears to induce the production of higher levels of the reactant (citrate) and lower levels of the product (Acetyl-CoA) (30). Upon VACV infection, we did not observe a significant increase in the protein levels of the mitochondrial citrate transporter (SLC25A1) or any obvious increases in levels of phosphorylated ACC1, which catalyzes the irreversible carboxylation of Acetyl-CoA required for fatty synthesis (**Fig S1**). Furthermore, an overall decrease in the steady-state levels of long-chain fatty acids was observed following VACV infection (30). Long-chain fatty acids are acylated and then carnitylated by carnitine palmitoyltransferase (CPT1), and are then transported into the mitochondrial matrix, where they undergo β-oxidation to fuel the TCA cycle (61). We have previously shown that the levels of carnitylated fatty acids increase following VACV infection, and inhibition of fatty acid β-oxidation suppresses VACV replication indicating an important role of VACV-induced upregulation of fatty acid β-oxidation in a VGF-dependent manner (30). It is known that ACLY positively regulates the carnitine system (53). Our findings indicate that the observed increase in ACLY phosphorylation in WT VACV infected cells (**Fig 1B, 1C, 3A)** led to increased neutral lipid droplets formation (**Fig 5A, 5B)**, geared toward generating β-oxidation intermediates (30) that are eventually recycled to the TCA cycle to generate energy (30, 58, 60).

**Figure S1.**
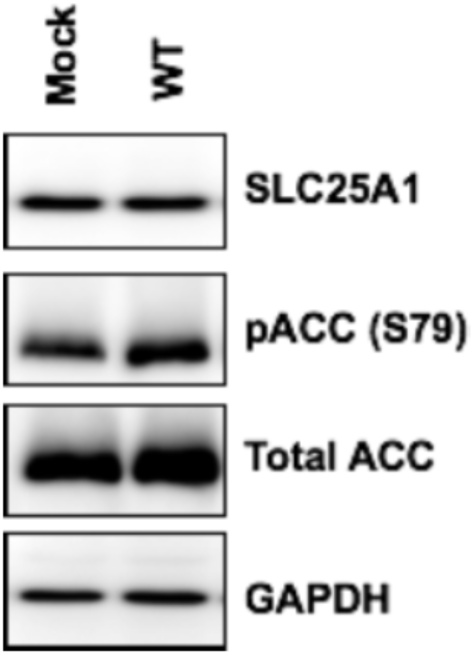
VACV infection does not induce the protein levels of mitochondrial citrate transporter or acetyl-CoA-CoA Carboxylase. HFFs were infected with MOI-5 of WT VACV. Western blotting analysis was performed to measure the levels of indicated proteins at 4 h post-infection (hpi).

VACV increases the levels of neutral lipid droplets in a VGF-dependent manner in HFFs. The production of lipid material in the cell relies on fatty acid synthesis, which is crucial for meeting various cellular requirements, including β-oxidation and increased membrane production. At the heart of this process is the synthesis of palmitate from acetyl-CoA and malonyl-CoA, which is facilitated by FASN. It has previously been shown in BSC-1 cells that palmitate is important during VACV replication to generate an energetically favorable environment through β-oxidation (58). Interestingly, our metabolic profiling of VACV-infected HFFs showed a significant decrease in the levels of both acetyl-CoA and palmitate (30). Our current findings that VACV infection induces the formation of lipid droplets in a VGF-dependent manner (**Fig 5A, 5B)**, provide a plausible explanation for these seemingly paradoxical findings.

While lipid droplets have been observed primarily in macrophage models following bacterial infection, the potential of viral infections in cells to elicit a similar response has not been extensively investigated. A limited number of studies have shown that viral infection of the positive-stranded RNA viruses, Sindbis and dengue virus, induces lipid droplet formation in the mosquito midgut cells (62). In another study, infection of mammalian cells with herpes simplex virus-1, influenza A virus, dengue virus, and zika virus showed transient induction of lipid droplets early during virus replication that corresponded with the detection of intracellular dsRNA and dsDNA (63). Interestingly, this induction of lipid droplets is independent of type-I interferon (IFN) and dependent on the EGFR-PI3K pathway. Because VACV infection produces dsRNA during the transcription of viral intermediate or late genes (64), it would be interesting to examine if the VGF-induced EGFR-PI3K pathway and dsRNA work in conjunction to enhance lipid droplet formation during VACV replication. Furthermore, a recent report indicates ISG15, an interferon-stimulated gene, is required for inducing lipid droplet formation during VACV infection of mouse bone marrow-derived macrophages (BMDM) (50). It is important to note that VACV encodes dozens of immune regulators and efficiently blocks interferon responses during productive infection of many cell types including HFFs (65–67). It remains to be tested if ISG15 is also involved in the VGF-EGFR-ALCY signaling cascade to induce fatty acid biosynthesis, and if so, functioning upstream or downstream of ACLY.

Our results raise an intriguing question: why does VACV go through such lengths to upregulate the levels of TCA cycle intermediates and the neutral lipid droplets? By upregulating ACLY phosphorylation and redirecting the host metabolism toward neutral lipid droplet formation, VACV could achieve multiple goals. First, because VACV is an enveloped virus, it requires lipid molecules to synthesize its membrane (68). It is possible that the lipids derived from the envelope of neutral lipid droplet levels may be used during virion morphogenesis. Second, the fatty acids derived from the neutral lipid droplets could provide the essential intermediates to generate β-oxidation substrates for maintaining an energy-rich state to support the increased demands associated with viral replication (58). Third, as seen in other viruses (63), lipid droplets could play vital roles in facilitating the magnitude of the early antiviral immune response during VACV infection. Because lipid droplets are highly dynamic organelles, further studies are warranted to decipher the exact functions of VACV-induced lipid droplets in virus replication.

VGF is crucial for the induction of ACLY phosphorylation in VACV-infected cells. VGF is a VACV protein that is secreted early following VACV infection and represents a viral homolog of cellular EGF and transforming growth factor (25, 27, 69). VGF is vital for the replication and virulence of VACV in animal models and quiescent cells (28, 29). Furthermore, VGF induces proliferative effects in infected cells and facilitates cellular motility and spread (22, 24) through the activation of the EGFR signaling cascade (22, 26). VACV activates the PI3K-Akt pathway early during infection (32), and Akt phosphorylation increases upon VACV infection in a VGF-dependent manner (22). Here, we showed that VACV induces ACLY phosphorylation via the VGF-induced EGFR-Akt signaling pathway starting early during infection demonstrating a crucial role for the functional interaction between VGF and ACLY in VACV replication. Taken together, these results combined with the report that VGF upregulates non-canonical STAT3 phosphorylation to induce citrate levels, our current study highlights the importance of VGF as a “master” regulator of cellular metabolic alterations during VACV infection.

ACLY is not the sole source of acetyl-CoA and is not exclusively localized to the cytosol (70). During nutrient-restricted conditions, such as starvation, the enzyme ACCS2 can convert acetate into acetyl-CoA (71, 72). During human cytomegalovirus (HCMV) infection, the loss of the ability to utilize citrate for Acetyl-CoA synthesis through ACLY has little effect on either lipid synthesis or viral growth because ACCS2 compensates for the loss of ACLY (73). Because acetate supplementation did not enhance VACV replication (**Fig S2**) and ACLY inhibition severely suppressed viral replication, the function of ACCS2 appears unlikely to be of similar importance as ACLY during VACV infection. Although ACLY is a predominantly cytosolic enzyme, several studies have reported its localization to the nucleus (70, 74). The acetyl-CoA generated in the nucleus by nuclear ACLY is vital for homologous recombination (74) and histone acetylation (70). Further studies remain necessary to determine the intracellular distribution of ACLY during VACV infection and the effects, if any, of altered localization patterns on the modulation of transcription.

**Figure S2.**
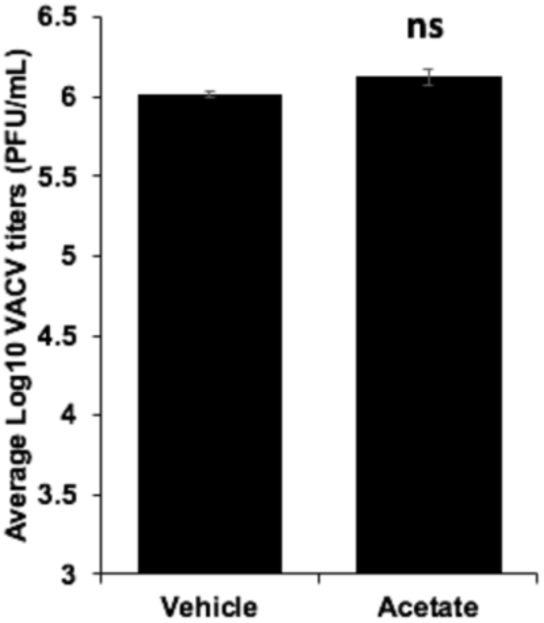
Supplementation of acetate does not enhance VACV replication. HFFs were infected with MOI-2 of WT VACV in the presence or absence of 5 mM Sodium acetate. VACV titers were determined by a plaque assay at 24 hpi.

ACLY phosphorylation during VACV infection occurs independently of EGFR-induced MAPK. The inhibition of EGFR, but not MAPK or STAT3, suppressed ACLY phosphorylation and Akt phosphorylation in VACV-infected cells but not in uninfected controls. Pioneering studies in the field have reported that ACLY phosphorylation is directly regulated by the PI3K-Akt pathway and the mitogen-activated protein kinase (MAPK) levels were not correlated with ACLY phosphorylation in lung adenocarcinoma cell lines (15), adipocytes (21), which is further supported by our findings in VACV infection.

Our current understanding of the role of ACLY in virus replication is limited. While chemical and genetic inhibition of ACLY suppressed the replication of SARS-CoV-2 (75), indicating it could be an important host factor governing the replication of the virus, it is yet unclear if the virus increases the phosphorylation or activity of ACLY. Another study in transformed HCC cells and transgenic mice expressing the Hepatitis B virus pre-S2 mutant in the liver showed increased ACLY phosphorylation through mTOR signaling to induce the levels of neutral lipids such as triglycerides and cholesterol (76), without showing direct effect of HBV infection on ACLY levels. In this regard, our finding sets a precedent to explore the effect of other viruses in modulating this critical host enzyme and how it affects their metabolism and replication. This will also open new avenues to develop novel targets for antiviral therapy.

In conclusion, this study demonstrated that the VGF-EGFR-Akt-induced ACLY phosphorylation and the synthesis of lipid droplet is crucial for VACV replication. Because poxviruses are widely used to develop oncolytic agents (3), and the ACLY-induced metabolism is often dysregulated in cancer cells (13, 20), our findings could lead to improvements in poxvirus-based oncolytic virotherapy and the development of better antipoxvirus agents.

## Acknowledgments

We thank Dr. Nicholas Wallace, Dr. Annie Newell-Fugate, and Dr. Bernard Moss for providing various cells and reagents. The work was supported by the National Institutes of Health (AI143709) to ZY and in part by a Graduate Student Summer Stipend to AP from the Johnson Cancer Research Center at Kansas State University. We thank Dr. Robert Burghardt and the image analysis lab (RRID: SCR_022479) at the School of Veterinary Medicine and Biomedical Sciences at Texas A&M University for providing the confocal microscopy facility.

